# Transgenic pyrimethamine resistant *Plasmodium berghei* as a model for *in vivo* anti-DHFR drug testing

**DOI:** 10.1101/2020.09.30.281055

**Authors:** Pongpisid Koonyosying, Natapong Jupatanakul, Jarunee Vanichtanankul, Thanaya Saeyang, Chatpong Pethrak, Jutharat Pengon, Wachiraporn Tipsuwan, Yongyuth Yuthavong, Sumalee Kamchonwongpaisan, Chairat Uthaipibull

## Abstract

Inhibitors for *Plasmodium falciparum* dihydrofolate reductase (DHFR) form an important class of antimalarial drugs widely used for malaria treatment, but have been compromised by development of resistance to the drugs. Mutations in DHFR are the main contributing factors to the resistance. Although new, rationally designed antifolates active against resistant *P. falciparum*, such as P218, have been developed, the activity against the quadruple mutant *P. falciparum* (V1/S) has only been demonstrated *in vitro*, and *in vivo* activity has only been shown in SCID mice. A convenient *in vivo* model for antifolate testing is desirable. In this study, the endogenous *P. berghei* dihydrofolate reductase-thymidylate synthase (*Pbdhfr*-ts) gene was successfully replaced by quadruple *dhfr-ts* mutant gene from *P. falciparum* (N51I+C59R+S108N+I164L). The transgenic parasite gained resistance to pyrimethamine but not to other class of antimalarial drugs. While 30 mg/kg of pyrimethamine could not inhibit the transgenic parasite, P218 could inhibit the transgenic parasite with the ED_50_ of 0.11±0.02 mg/kg, a level similar to the *P. falciparum* in SCID mice model. These results demonstrated the validity of our model and showed that P218 was very potent against quadruple *Pfdhfr-ts* mutant parasite, *in vivo*.

## 1. Introduction

Malaria continues to inflict extensive morbidity and mortality in many endemic areas of the world (World Health Organization, 2019). This mosquito-borne disease is caused by parasites in the genus *Plasmodium* with *Plasmodium falciparum* causing the most severe malaria (WHO, 2014). With effective vaccine still under development, antimalarial drugs play an important role to reduce burden of malaria in humans.

*P. falciparum* dihydrofolate reductase-thymidylate synthase (PfDHFR-TS) is an enzyme involved in folate metabolism and DNA synthesis thus essential for this highly replicative parasite. Selective inhibitors for PfDHFR-TS such as cycloguanil or pyrimethamine were widely used for malaria treatment and control, but resistance emerged soon after introduction of the drugs. Mutations on the *dhfr-ts* are the major contribution to the resistance to these DHFR inhibitors (Gregson and Plowe, 2005; Müller and Hyde, 2013; Yuthavong et al., 2006). Pyrimethamine resistance from *dhfr* mutations develops progressively in *P. falciparum*, and quadruple mutant (N51I+C59R+S108N+I164L) leads to complete treatment failure of the drug (Ahmed et al., 2006; Sirawaraporn, 1998).

Availability of crystal structures of these mutated enzymes allows rational development of new anti-PfDHFR agents that can inhibit quadruple *dhfr* mutant *P. falciparum* as well as wild-type and other mutant forms, with P218 as a lead compound (Yuthavong et al., 2012). During preclinical development of new drug candidates, data of *in vivo* efficacy of the drug is needed. Although the *P. falciparum* infection in severe combined immunodeficient (SCID) mice model is available, a more convenient model is desirable. *Plasmodium berghei* has long been used as a model organism for *in vivo* antimalarial drug development (Goodwin, 1949; Jiménez-Díaz et al., 2013). Although pyrimethamine resistant *P. berghei* and *P. chabaudi* strains were developed through drug pressure (Hayton et al., 2002; Nuralitha et al., 2016; van Dijk et al., 1994), these parasites might not be optimal proxies for anti-DHFR-TS compounds structurally designed to inhibit quadruple mutant PfDHFR-TS.

To facilitate preclinical development of anti-DHFR compounds against pyrimethamine resistant parasite, the wild-type *P. berghei dhfr-ts* gene (*Pbdhfr-ts*) was replaced with quadruple mutant *Pfdhfr-ts* through homologous recombination. The transgenic parasite was then used to demonstrate *in vivo* efficacy of P218 against pyrimethamine resistant parasite.

## 2. Materials and methods

### 2.1 Ethic statement

This study was strictly carried out in accordance to the recommendations in the Guide for the Care and Use of Laboratory Animals of the National Institutes of Health, and Thailand’s National research council. The animal protocol was approved by the BIOTEC Institutional Animal Care and Use Committee (Permit number: BT-Animal 27/2562).

### 2.2 Animal and Parasite strains

Eight weeks old female ICR mice were used for all the experiments in this study. The mice were obtained from the National Laboratory Animal Center, Mahidol University, Thailand. The mice were acclimatized for at least three days before experiments and were maintained at 22±2 °C. Water and food were provided *ad libitum. P. berghei* ANKA 676m1cl1, expressing GFP-luciferase (MRA-868) was used as a parental strain to generate transgenic *P. berghei*.

### 2.3 Transfection plasmid construction

pY003V1S plasmid for *P. berghei* transfection was constructed to contain *Pfdhfr-ts* with amino acid mutations in *dhfr* at the positions N51I, C59R, S108N and I164L, the same as the V1/S strain *P. falciparum*, flanking by 5’ and 3’ untranslated region (UTR) of *Pbdhfr-ts* (Figure 1A). The quadruple mutant *Pfdhfr* fragment was amplified from pCas.SgDHFR.HR.V1S plasmid (Posayapisit et al., 2020) and cloned into pY003 plasmid at *Nco*I and *Spe*I restriction sites. The *P. berghei* UTRs serve as sites for double crossover homologous recombination at the endogenous *Pbdhfr-ts* locus. To generate integrative transformant, the pY003V1S plasmid was linearized at the *Hind*III*, Kas*I *and Sca*I site, before transfection experiments.

**Figure 1.**
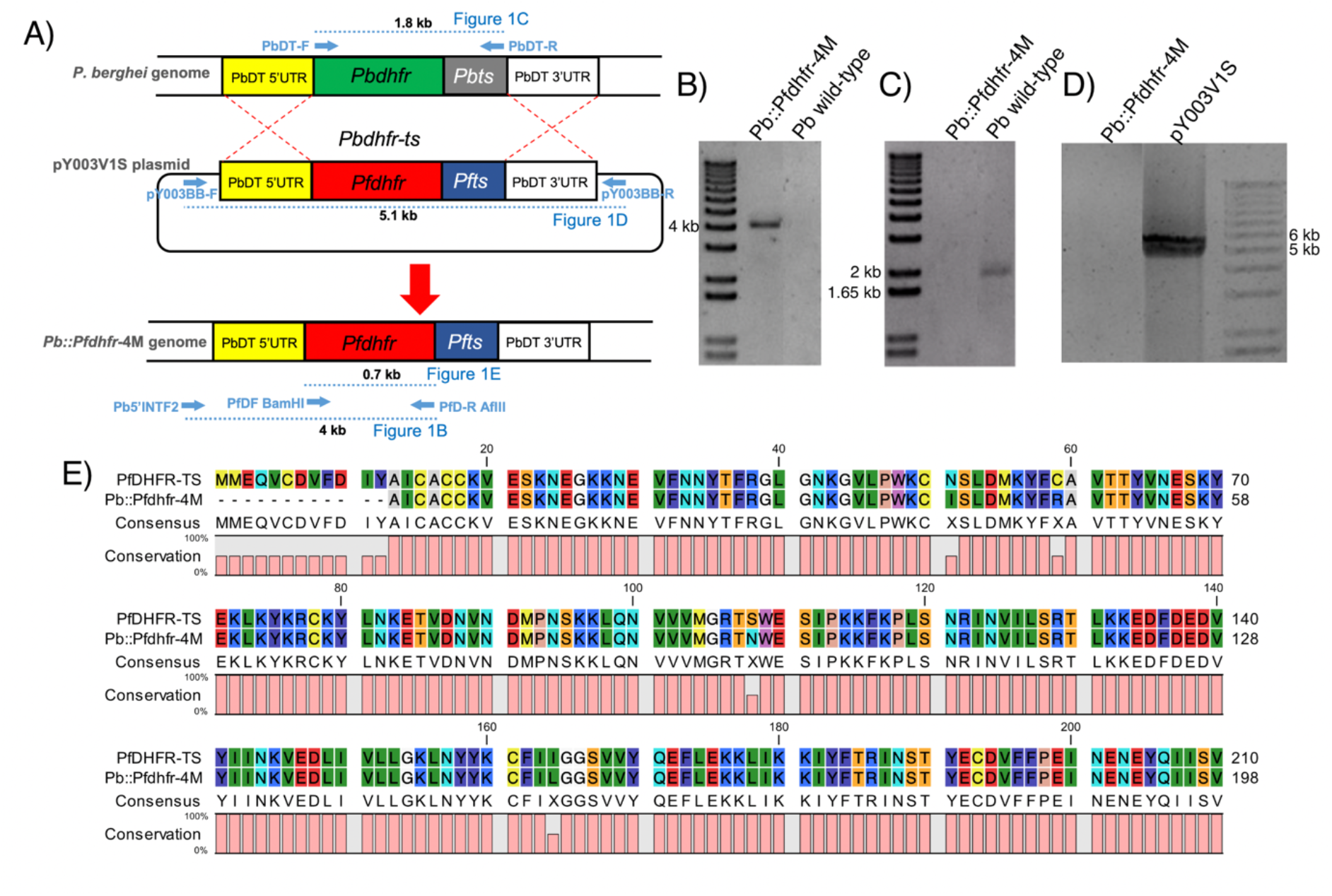
Construction of *Pb::Pfdhfr*-4M transgenic parasite. A) schematic diagram of homologous recombination of quadruple mutant *Pfdhfr* into *P. berghei* genome. Dotted blue lines represent the amplicons from PCR characterization. B) 4 kb amplicon from 5’PbINTF2 and PfDR AflII primers. C) 1.8 kb amplicon from PbDTF and PbDTR primers. D) 5.1 kb amplicon from pY003BBF and pY003BBR primers. E) alignment of translated amino acid sequence between the wild-type *Pfdhfr* to the *Pfdhfr* gene from *Pb::Pfdhfr*-4M transgenic parasite that has mutations at N51I, C59R, S108N, and I164L. The *Pfdhfr* fragment was amplified and sequenced with PfDF BamHI and PfDR AflII primers.

### 2.4 P. berghei transfection

*P. berghei* was transfected following a published protocol (Franke-Fayard et al., 2004). Briefly, heparinized *P. berghei* infected blood was diluted to 10^6^ infected red blood cells per μl with RPMI-1640 medium supplemented with 0.2% NaHCO_3_, 30% FBS, 200 U/ml Heparin and 50 μg/ml Neomycin sulfate, filled with 94% N_2_, 1% O_2_, 5% CO_2_, then cultured overnight at 37°C, with shaking at 90 rounds per minute. *P. berghei* schizonts were purified on a Nycodenz gradient (Alere Technologies AS, Norway), washed, pelleted by centrifugation, then 10^6^ purified schizonts were resuspended in Nucleofector solution (Lonza, NJ, USA) containing 10 μg of linearized pY003V1S plasmid, and electroporated with program U33 using the Amaxa Nucleofector device (Lonza, NJ, USA). Transfected parasites were then injected intravenously into naïve ICR mice and selected by pyrimethamine administered in the drinking water (120 μg/ml). Transfected parasites were cloned by limiting dilution.

### 2.5 Characterization of the transgenic parasite

Genomic DNA of the transfected parasite was used to confirm quadruple mutant *Pfdhfr-ts* integration by PCR (Figure 1A). PbDTF (5’ GGGGGGGGCATATGGAAGACTTATCTGAAACATTCG 3’) and PbDTR (5’GGACTAGTTTAAGCTGCCATATCC 3’) were used to confirm the absence of endogenous *Pbdhfr-ts*. pY003BBF (5’ GCTATGACCATGATTACGCCAAGC 3’) and pY003BBR (5’CAGATTGTACTGAGAGTGCACCATATGC 3’) were used to confirm the absence of the pY003V1S plasmid in episomal form. Primers 5’PbINTF2 (5’ GCTACATAACTTCCATACATGTCTGGGC 3’) and PfDR AflII (5’CTTTGTCATCATTCTTAAGAGGC 3’) were used to confirm *Pfdhfr-ts* integration.

To confirm the sequence of quadruple mutant *Pfdhfr-ts* in the transgenic parasite, *Pfdhfr* gene fragment was amplified by PCR using PfDF BamHI (5’CGGTGGATCCATGATGGAACAAG 3’) and PfDR AflII. The amplicon was then used for sequencing by the same primers. Multiple sequence alignment was performed using CLC sequence viewer (QIAGEN Aarhus A/S version 7.6.1).

### 2.6 In vivo antimalarial drug testing

The 4-day suppressive test (Peters et al., 1975) was used for *in vivo* antimalarial drug testing. Briefly, 8-week-old ICR mice were intraperitonially injected with 10^7^ infected erythrocytes of either the wild-type *P. berghei* ANKA or the transgenic parasite. Antimalarial compounds were freshly prepared in suspension vehicle (20% DMSO, 7% Tween-80, 3% Ethanol) then orally administered at 6, 24, 48, and 72 hours after parasite inoculation. The parasitemia in mice after 4-day suppressive test was determined with Giemsa stained thin blood smear. Pyrimethamine (Sigma, Cat# P7771) and chloroquine diphosphate salt (Sigma, Cat# C6628) were used as standard antimalarial drugs.

### 2.7 Statistical analyses

Dose response analyses were performed using 4-parameter log-logistic regression model using the *drc* package (Ritz et al., 2015) in the R program (R Core Team, 2020).

## 3. Results

### 3.1 Generation of transgenic P. berghei stably expressing quadruple mutant PfDHFR-TS

*P. berghei* ANKA 676m1cl1 expressing GFP-luciferase was successfully transfected with pY003V1S plasmid to replace wild-type *Pbdhfr-ts* with quadruple mutant *Pfdhfr-ts*. After positive selection with PYR and cloning, the transgenic parasite *Pb::Pfdhfr*-4M was obtained. The presence of *Pfdhfr-ts* gene in the transgenic parasite was confirmed by PCR (Figure 2). The transgenic parasite has quadruple mutant *Pfdhfr-ts* integrated into the genome (Figure 2B), has *Pbdhfr-ts* removed (Figure 2C), and does not have episomal pY003V1S plasmid (Figure 2D). Sequencing results confirmed correct sequence of quadruple mutant *Pfdhfr-ts* in the transgenic parasite with mutations at four amino acid positions N51I, C59R, S108N, and I164L (Figure 2E).

**Figure 2.**
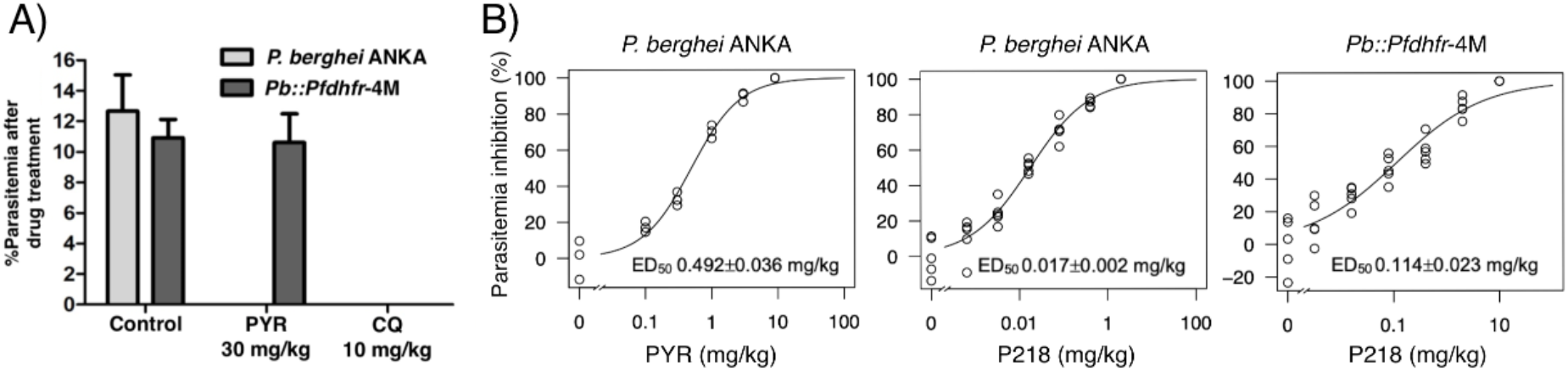
Transgenic *Pb::Pfdhfr*-4M gained pyrimethamine (PYR) resistance but P218 was still highly potent against the parasite. A) Bar chart representing parasitemia of mice after oral antimalarial drug treatment with 4-day suppressive test. Error bars represent SEM. Vehicle without drug was given in the control group. CQ: Chloroquine. B) Dose response curves of pyrimethamine and P218 in the wild-type parasite, and P218 in *Pb::Pfdhfr*-4M. Y-axis represent percent inhibition of parasitemia after 4-day suppressive test compared to the average of untreated control group.

### 3.2 Pb::Pfdhfr-4M gain resistance to pyrimethamine

While pyrimethamine at a dose of 30 mg/kg could completely inhibit the growth of parental *P. berghei* ANKA, it could not inhibit the *Pb::Pfdhfr*-4M parasite. After 4 days of pyrimethamine administration, parasitemia of the transgenic parasite was 10.62±1.87%, which is similar to the control group (Figure 2A). In addition to the antifolate, chloroquine was used as a non-antifolate drug control. Both the transgenic *Pb::Pfdhfr*-4M and wild-type parasites were susceptible to chloroquine at 10 mg/kg (Figure 2A). These results demonstrated that the transgenic parasite gained pyrimethamine resistance but not another class of antimalarial compounds.

### 3.3 P218 was highly potent against both the wild-type and Pb::Pfdhfr-4M in vivo

The dose response analyses revealed that *in vivo* ED_50_ of P218 was 0.017±0.002 mg/kg and 0.114±0.023 mg/kg in the wild-type *P. berghei* and *Pb::Pfdhfr*-4M, respectively (Figure 2B). Although the quadruple *dhfr* mutation increased *in vivo* ED_50_ of P218 by 6.7 times, the ED_50_ of P218 against the *Pb::Pfdhfr*-4M was still 4.32 times lower than ED_50_ of pyrimethamine against the wild-type parasite, which was 0.492±0.036 mg/kg (Figure 2B).

## 4. Discussion

Up to now, there has been a lack of a cheap and reliable *in vivo* model to test anti-DHFR compound that designed specifically to target pyrimethamine resistant *P. falciparum* parasites. The pyrimethamine resistant *P. berghei* and *P. chabaudi* generated from drug selection pressure (Hayton et al., 2002; Nuralitha et al., 2016; van Dijk et al., 1994) might not be good representatives for the pyrimethamine-resistant *P. falciparum* due to different binding affinity of the mutated *Pbdhfr* compared to the mutated *Pfdhfr*. Although it is possible to use *P. falciparum*-in a humanized mouse model to evaluate potency of antimalarial compounds *in vivo*, such model requires tremendous resources and availability of the humanized mice. This study presents the generation of transgenic *P. berghei* model with its anti-DHFR sensitive *dhfr-ts* gene replaced by pyrimethamine-resistant quadruple mutant from *P. falciparum*. The transgenic parasite is a reliable and cheap preclinical *in vivo* model to evaluate the potency of rationally designed *Pfdhfr*-specific inhibitors. Since this transgenic parasite does not require special handlings, routine *in vivo* evaluation of *Pfdhfr*-specific inhibitors is now possible in laboratories with basic animal facility.

The transgenic parasite was validated by testing *in vivo* efficacy of P218, a novel antifolate compound rationally designed to inhibit pyrimethamine resistant quadruple *dhfr* mutant *P. falciparum* (Yuthavong et al., 2012). Our study demonstrated lower ED_50_ of P218 against the quadruple *Pfdhfr* mutant parasite compared to the ED_50_ of pyrimethamine in the wild-type parasite. The ED_50_ obtained from the *Pb::Pfdhfr*-4M model was similar to the previously published ED_50_ of 0.3 mg/kg against V1/S strain *P. falciparum* in the SCID mice (Yuthavong et al., 2012), validating that our transgenic parasite is a cheaper and more convenient alternative preclinical model for *in vivo* anti-DHFR testing. Together with a good safety profile shown in the first-in human clinical trial (Chughlay et al., 2020), these results support that P218 will be highly effective for malaria treatment even in the area with predominant pyrimethamine-resistant parasite and an invaluable tool for malaria elimination.

## Acknowledgement

This work was financially supported by UNICEF/UNDP/World Bank/WHO Special Program for Research and Training in Tropical Diseases (TDR) grant (#A60924) to CU, Research Chair Grant (P-1850116) from the National Science and Technology Development Agency (NSTDA), Thailand to SK, and BIOTEC Research unit director initiative grant to NJ (P-1851424). P218 compound was kindly provided by Medicine for Malaria Venture (MMV). *P. berghei* ANKA 676m1cl1 expressing GFP-luciferase (MRA-868), contributed by Chris J. Janse and Andrew P. Waters was obtained through BEI Resources, NIAID, NIH.

## Author contributions

**Pongpisid Koonyosying:** Methodology, Validation, Formal analysis, Investigation, Writing - Original Draft

**Natapong Jupatanakul:** Conceptualization, Methodology, Validation, Formal analysis, Investigation, Writing - Original Draft, Writing - Review & Editing, Visualization, Supervision, Project administration, Funding acquisition

**Jarunee Vanichtanankul:** Methodology, Investigation

**Thanaya Saeyang:** Investigation

**Chatpong Pethrak:** Methodology, Investigation

**Jutharat Pengon:** Methodology, Investigation

**Wachiraporn Tipsuwan**: Methodology, Investigation

**Yongyuth Yuthavong:** Resources, Writing - Review & Editing

**Sumalee Kamchonwongpaisan:** Methodology, Resources, Funding acquisition, Writing - Review & Editing, Funding acquisition

**Chairat Uthaipibull:** Conceptualization, Methodology, Resources, Writing - Review & Editing, Supervision, Funding acquisition

## References

Ahmed, A., Das, M.K., Dev, V., Saifi, M.A., Wajihullah, Sharma, Y.D., 2006. Quadruple Mutations in Dihydrofolate Reductase of *Plasmodium falciparum* Isolates from Car Nicobar Island, India. Antimicrobial Agents and Chemotherapy 50, 1546–1549. https://doi.org/10.1128/aac.50.4.1546-1549.2006

Chughlay, M.F., Rossignol, E., Donini, C., Gaaloul, M.E., Lorch, U., Coates, S., Langdon, G., Hammond, T., Möhrle, J., Chalon, S., 2020. First-in-human clinical trial to assess the safety, tolerability and pharmacokinetics of P218, a novel candidate for malaria chemoprotection. British Journal of Clinical Pharmacology 86, 1113–1124. https://doi.org/10.1111/bcp.14219

Franke-Fayard, B., Trueman, H., Ramesar, J., Mendoza, J., van der Keur, M., van der Linden, R., Sinden, R.E., Waters, A.P., Janse, C.J., 2004. A *Plasmodium berghei* reference line that constitutively expresses GFP at a high level throughout the complete life cycle. Molecular and Biochemical Parasitology 137, 23–33. https://doi.org/10.1016/j.molbiopara.2004.04.007

Goodwin, L.G., 1949. Response of *Plasmodium berghei* to Anti-malarial Drugs. Nature 164, 1133–1133. https://doi.org/10.1038/1641133a0

Gregson, A., Plowe, C.V., 2005. Mechanisms of Resistance of Malaria Parasites to Antifolates. Pharmacol Rev 57, 117–145. https://doi.org/10.1124/pr.57.1.4

Hayton, K., Ranford-Cartwright, L.C., Walliker, D., 2002. Sulfadoxine-Pyrimethamine Resistance in the Rodent Malaria Parasite *Plasmodium chabaudi*. Antimicrobial Agents and Chemotherapy 46, 2482–2489. https://doi.org/10.1128/AAC.46.8.2482-2489.2002

Jiménez-Díaz, M.B., Viera, S., Ibáñez, J., Mulet, T., Magán-Marchal, N., Garuti, H., Gómez, V., Cortés-Gil, L., Martínez, A., Ferrer, S., Fraile, M.T., Calderón, F., Fernández, E., Shultz, L.D., Leroy, D., Wilson, D.M., García-Bustos, J.F., Gamo, F.J., Angulo-Barturen, I., 2013. A New In Vivo Screening Paradigm to Accelerate Antimalarial Drug Discovery. PLOS ONE 8, e66967. https://doi.org/10.1371/journal.pone.0066967

Müller, I.B., Hyde, J.E., 2013. Folate metabolism in human malaria parasites—75 years on. Molecular and Biochemical Parasitology 188, 63–77. https://doi.org/10.1016/j.molbiopara.2013.02.008

Nuralitha, S., Siregar, J.E., Syafruddin, D., Roelands, J., Verhoef, J., Hoepelman, A.I.M., Marzuki, S., 2016. Within-Host Selection of Drug Resistance in a Mouse Model of Repeated Incomplete Malaria Treatment: Comparison between Atovaquone and Pyrimethamine. Antimicrobial Agents and Chemotherapy 60, 258–263. https://doi.org/10.1128/AAC.00538-15

Peters, W., Portus, J.H., Robinson, B.L., 1975. The chemotherapy of rodent malaria, XXII: The value of drug-resistant strains of *P. berghei* in screening for blood schizontocidal activity. Annals of Tropical Medicine & Parasitology 69, 155–171. https://doi.org/10.1080/00034983.1975.11686997

Posayapisit, N., Pengon, J., Prommana, P., Shoram, M., Yuthavong, Y., Uthaipibull, C., Kamchonwongpaisan, S., Jupatanakul, N., 2020. Transgenic pyrimethamine-resistant *P. falciparum* reveals transmission blocking potency of P218, a novel antifolate. bioRxiv 2020.09.06.284786. https://doi.org/10.1101/2020.09.06.284786

R Core Team, 2020. R: The R Project for Statistical Computing. R Foundation for Statistical Computing, Vienna, Austria.

Ritz, C., Baty, F., Streibig, J.C., Gerhard, D., 2015. Dose-Response Analysis Using R 10, e0146021. https://doi.org/10.1371/journal.pone.0146021

Sirawaraporn, W., 1998. Dihydrofolate reductase and antifolate resistance in malaria. Drug Resistance Updates 1, 397–406. https://doi.org/10.1016/S1368-7646(98)80015-0

van Dijk, M.R., McConkey, G.A., Vinkenoog, R., Waters, A.P., Janse, C.J., 1994. Mechanisms of pyrimethamine resistance in two different strains of *Plasmodium berghei*. Molecular and Biochemical Parasitology 68, 167–171. https://doi.org/10.1016/0166-6851(94)00163-4

WHO, 2014. Severe Malaria. Trop Med Int Health 19, 7–131. https://doi.org/10.1111/tmi.12313_2

World Health Organization, 2019. World Malaria Report 2019. WORLD HEALTH ORGANIZATION, S.l.

Yuthavong, Y., Kamchonwongpaisan, S., Leartsakulpanich, U., Chitnumsub, P., 2006. Folate metabolism as a source of molecular targets for antimalarials. Future Microbiology 1, 113–125. https://doi.org/10.2217/17460913.1.1.113

Yuthavong, Y., Tarnchompoo, B., Vilaivan, T., Chitnumsub, P., Kamchonwongpaisan, S., Charman, S.A., McLennan, D.N., White, K.L., Vivas, L., Bongard, E., Thongphanchang, C., Taweechai, S., Vanichtanankul, J., Rattanajak, R., Arwon, U., Fantauzzi, P., Yuvaniyama, J., Charman, W.N., Matthews, D., 2012. Malarial dihydrofolate reductase as a paradigm for drug development against a resistance-compromised target. PNAS. https://doi.org/10.1073/pnas.1204556109

